# Enhancing Biomarker-Based Oncology Trial Matching Using Large Language Models

**DOI:** 10.1101/2024.09.13.612922

**Authors:** Nour Al Khoury, Maqsood Shaik, Ricardo Wurmus, Altuna Akalin

## Abstract

Clinical trials are an essential component of drug development for new cancer treatments, yet the information required to determine a patient’s eligibility for enrollment is scattered in large amounts of unstructured text. Genomic biomarkers are especially important in precision medicine and targeted therapies, making them essential for matching patients to appropriate trials. Large language models (LLMs) offer a promising solution for extracting this information from clinical trial data, aiding both physicians and patients in identifying suitable matches. In this study, we explore various LLM strategies for extracting genetic biomarkers from oncology trials to improve patient enrollment rates. Our results show that open-source language models, when applied out-of-the-box, effectively capture complex logical expressions and structure genomic biomarkers in disjunctive normal form, outperforming closed-source models such as GPT-4 and GPT-3.5-Turbo. Furthermore, fine-tuning these open-source models with additional data significantly enhances their performance.

## Introduction

Cancer is one of the leading causes of death in the world ^1^. According to the Global Cancer Observatory (GLOBOCAN) statistics produced by the International Agency for Research on Cancer (IARC), the estimation for the year 2022 is almost 20 million new cases of cancer and around 9.7 million cancer deaths, globally ^2^. Surgery, chemotherapy, and radiotherapy, individually or in combination, represent the standard treatments for cancer patients ^3,4,5^. Nevertheless, although these traditional methods are successful, they still have limitations. For instance, surgical removal of cancer is the most risk-averse option, but it is only possible in its early stages ^5^. Moreover, chemotherapy and radiotherapy are not effective for all cancer types ^3^, as they are not devoid of side effects since they attack cancerous cells as well as healthy cells ^4,6^. A promising direction in cancer treatment is precision medicine, which involves studying the patients’ genetics, lifestyle, and environmental information to guide treatment decisions ^7^. This approach improves care quality while reducing the need for unnecessary diagnostic tests and therapies by guiding healthcare decisions toward the most effective treatments for a given patient ^8^.

Therapies targeting specific genes or associated with certain response biomarkers have demonstrated considerable promise in the treatment of patients with different cancer types ^1,9,10^. According to a study conducted on clinical trial records from 2000 to 2015, clinical trials that use biomarkers for patient stratification have higher success rates, particularly for oncology clinical trials ^11^. As of 2022, the FDA acknowledges 45 genes and MSI-H and TMB-H as biomarkers that can predict a patient’s response to a drug and five tumor-agnostic genomic biomarkers ^12^. Despite the benefits of increasing patient participation in clinical trials, such as accelerating the development of treatments and improving trial generalizability ^13^, many clinical trials fail to be completed due to poor recruitment, amongst other reasons ^14,15,16^. Even though the final decision to participate in a trial is in the hands of the patient, various barriers often prevent patients from being offered the opportunity to enroll in the trial enrollment process ^17^. A systematic review defined four barrier domains affecting a patient’s participation in a trial post-diagnosis, leading to only 8.1% of 8883 patients being enrolled ^17^. Physicians may not discuss possible trials due to barriers such as treatment preferences ^17^, lack of awareness ^18^, or perceiving the process as time-consuming ^19,20,21^. Improving enrollment relies on effective methods to match patients to clinical trials. Although clinical trial information is available through platforms like clinicaltrials.gov ^17^, the lack of structured data and non-standard naming conventions, coupled with the increasing number of trials lead to a time-consuming and overwhelming search process. In oncology, matching a potential trial to a patient could be even more complicated due to the increasing use of biomarkers and non-standard naming conventions.

The issues outlined above create a need for tools that facilitate the search process, narrowing down to the candidate ongoing trials most suitable for patients, increasing the likelihood of their enrollment in a suitable trial. Therefore, natural language processing (NLP) has emerged to expedite the clinical trial matching process ^22^. Amidst the rise of LLMs and the large number of ongoing clinical trials, many approaches have emerged to automate patient-to-trial matching, speeding up the process and increasing the chances of finding a suitable trial. Existing approaches apply either an end-to-end matching ^23,24,25^ or a structure-then-match strategy ^22,26^. In an end-to-end strategy, the LLM is used to compare a patient’s record with an unstructured clinical trial document and return a final decision on the trial’s eligibility or ranking. In the structure-then-match strategy, the LLM is used to extract and structure entities from the clinical trial text, followed by the matching process. This process is straightforward, assuming the structuring part is done well.

In this study, we took a structure-then-match approach to enhance biomarker-based patient-to-trial matching by improving the biomarker extraction process from clinical trials. We evaluate the model’s ability to simultaneously extract genomic biomarkers from a clinical trial, pre-process the extracted data, and structure the logical connections (AND/OR) between biomarkers in the disjunctive normal form (DNF) ^22^. The DNF is a representation of a logical formula as a disjunction (OR) of conjunctions (ANDs).

To achieve this, we investigated multiple large language models using various prompting techniques and fine-tuned an open-source model as depicted in Figure 1.

**Figure 1:**
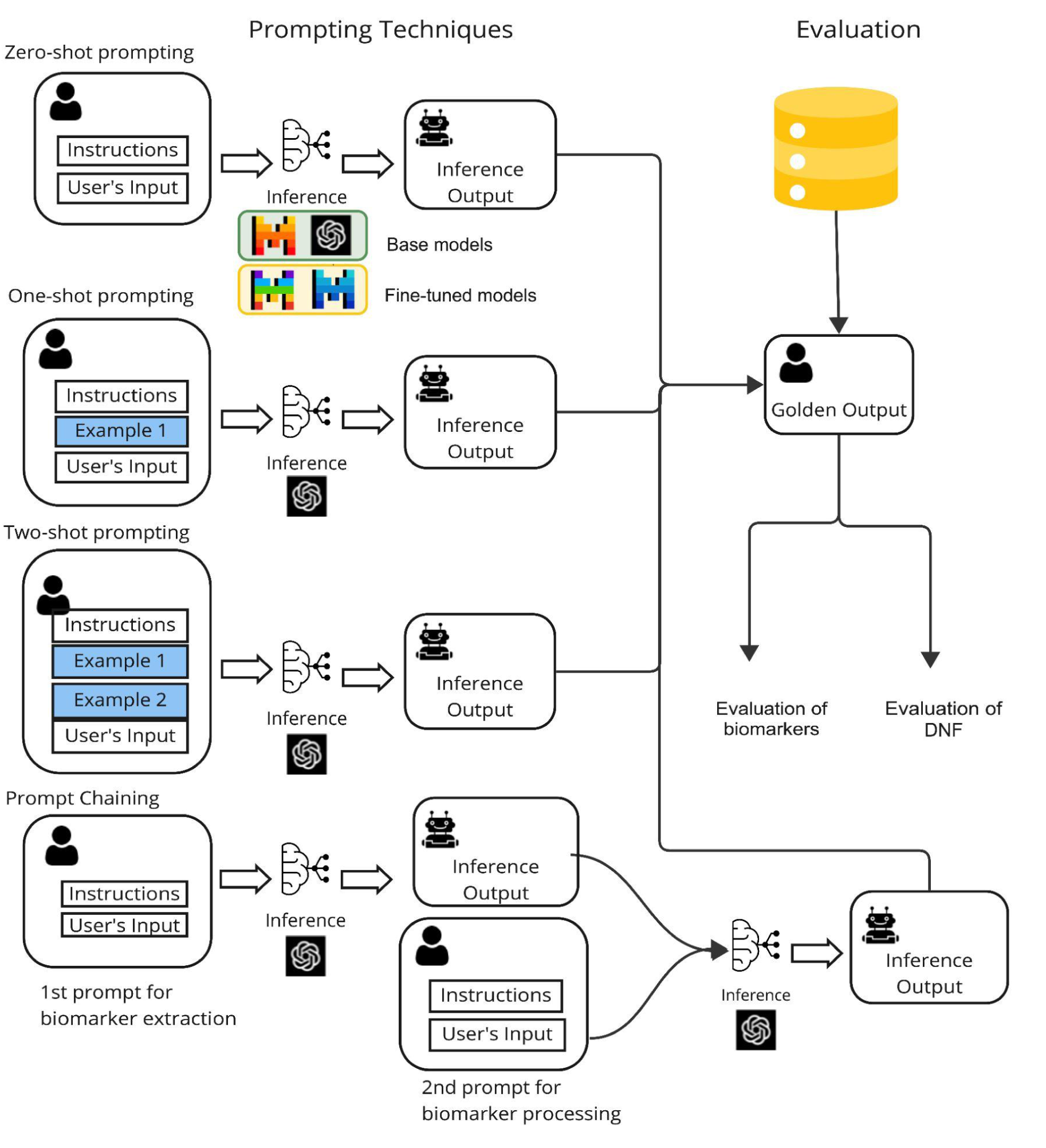
Overall Framework for Biomarker Extraction and Evaluation. This figure illustrates the prompting techniques explored for extracting genetic biomarkers from clinical trials. It underlines the evaluation of the extraction process both with and without assessing the model’s capability to organize the biomarkers in Disjunctive Normal Form (DNF). The framework includes several prompting approaches, including zero-shot, one-shot, two-shot prompting, and prompt chaining, which involves sequential prompts for biomarker extraction and processing. The framework also highlights the usage of fine-tuned models.

With our fine-tuned model, we achieved superior performance in extracting and structuring biomarkers in the DNF form.

## Results

### Data curation and trial data characteristics

As biomarkers are an important part of cancer drug development and diagnostics, we focused on curating a representative data set. The CIViC database (https://civicdb.org) is an open-source knowledgebase that includes extensive cancer-related biomarker datasets. We identified 500 biomarkers related to cancer diagnostics and therapy.

Based on the large-scale AACR-genie cancer patient cohort, which included data from 171,957 patients, we estimate that at least a subset of these biomarkers are present in 23.56% of cancer patients. In terms of cancer types, colorectal cancer, breast cancer, and glioma are those with most frequent biomarkers (Figure 2)

**Figure 2:**
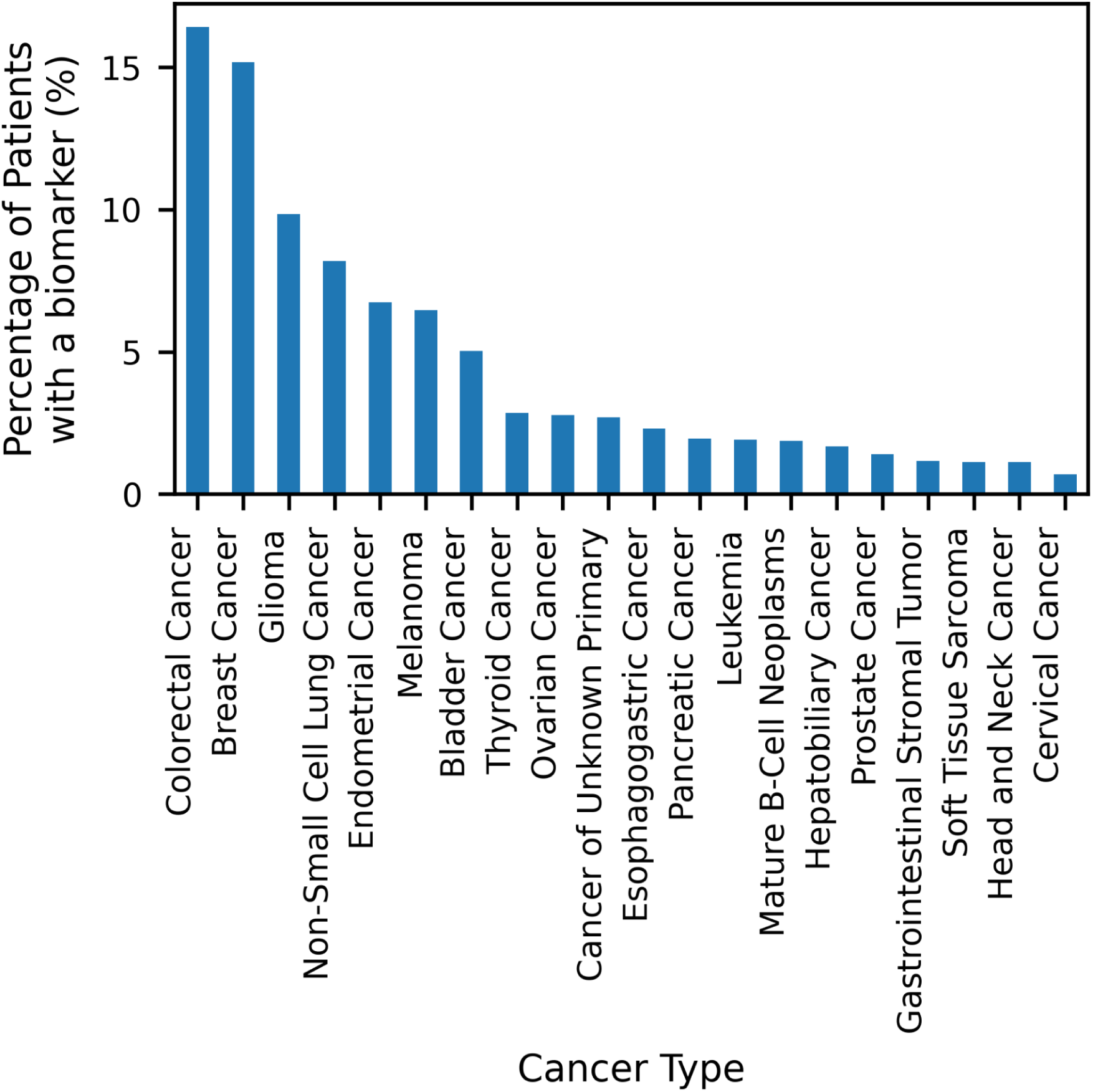
Distribution of Top 20 Cancer Types. Bar chart showing the distribution of the top 20 cancer types among patients who matched at least one of the 500 biomarkers selected from CIViC. The patients’ records are collected from the American Association for Cancer Research (AACR) Project GENIE.

We have obtained the ongoing oncology clinical trials from clinicaltrials.gov. A clinical trial typically includes a summary of the trial plan, followed by a detailed description, and the inclusion and exclusion eligibility criteria required for enrollment. Then we query the trials with the list of biomarkers from the CIViC database. This allowed us to select 296 unique trials with the potential presence of a biomarker in the eligibility criteria. From these trials, we manually annotated 166 of the trials, detailing the inclusion and exclusion biomarkers for each trial in JSON format. We removed one outlier sample that had a significantly larger token count (Supplementary Figure 1). We then split the data into a 70:30 ratio resulting in 116 training samples and 50 testing samples.

We prepared the training dataset for fine-tuning with Direct Preference Optimization (DPO) and split it into an 80:20 ratio which resulted in the first fine-tuning dataset DPO-92 with 92 samples as training set and 23 samples as validation set. To create a second fine-tuning dataset, the original dataset, DPO-92 was augmented with 80 synthetically generated samples using GPT-4 (See “Generation of Synthetic Dataset” for details). These additional samples underwent the same process of preparation for fine-tuning with DPO. The combined dataset was split following an 80:20 ratio resulting in the dataset DPO-156 with 156 samples for training and 39 samples for validation. Refer to “DPO Dataset Preparation” for more details.

### Zero-shot, Few-shot and Prompt Chaining Performance

We employ the models using three prompting techniques: zero-shot prompting where the model is given instructions to be followed, prompt chaining where the task is performed using a chain of requests to the model, and finally, few-shot prompting where the model is provided with examples to demonstrate the task. Refer to the methods section for a detailed description of the prompting techniques.

In Table 1 we report the models’ performance with the prompting techniques using a preliminary evaluation approach where we only measure the model’s ability to extract the inclusion and exclusion biomarkers found in the clinical trial. With zero-shot prompting, GPT-3.5-Turbo had a moderate performance in the extraction of the inclusion biomarkers with an F2 score 0.45. However, the model struggled with the extraction of exclusion biomarkers resulting in one of the lowest F2 scores (0.06).

**Table 1:**
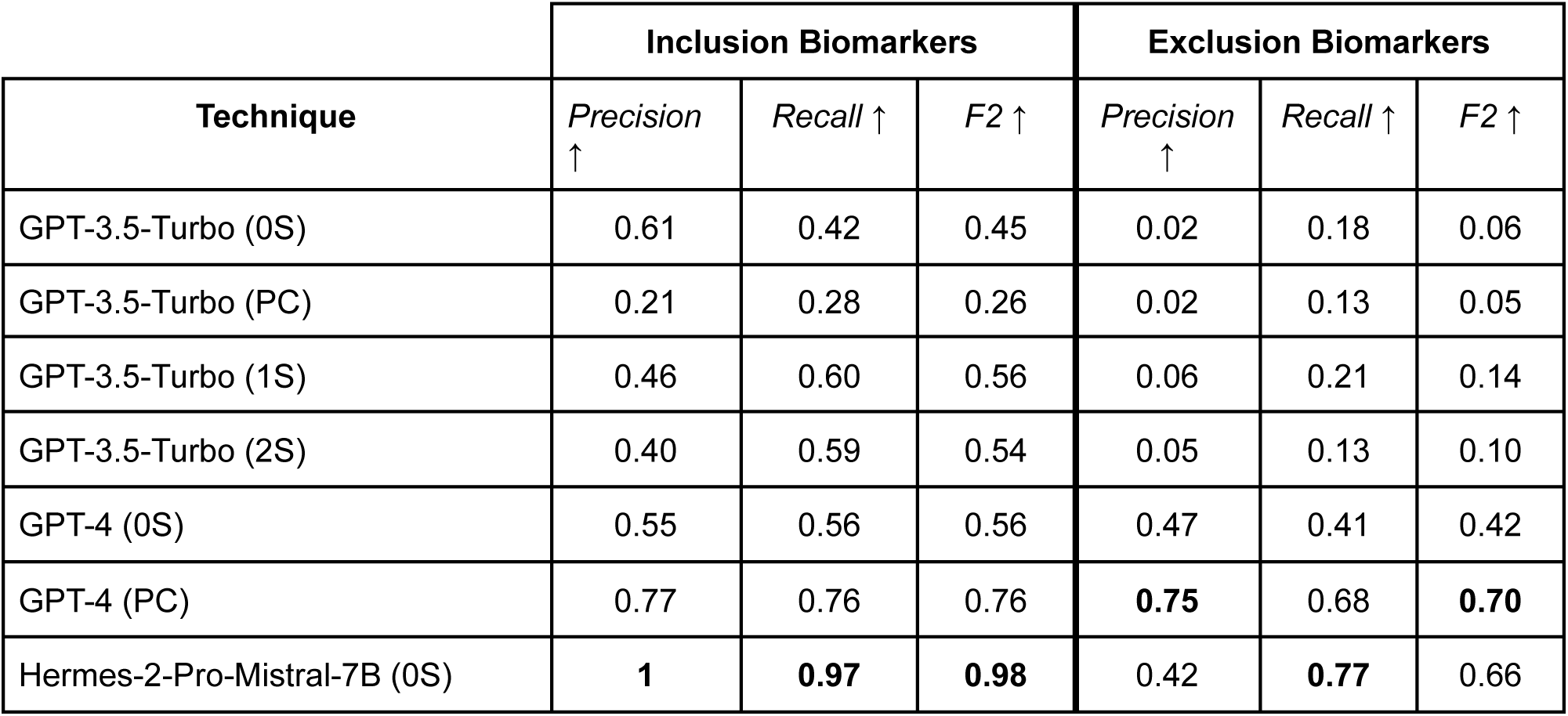
Performance of the open-source and closed-source base models using the different prompting techniques in extracting the inclusion and exclusion biomarkers from the clinical trials free-text documents. The prompting techniques applied include zero-shot (0S) prompting, where the prompt describes the input, output, and task; one-shot (1S) prompting, where an additional example is provided to demonstrate the task; two-shot (2S) prompting, where two examples are given to illustrate the task; and prompt chaining (PC), which divides the task into subtasks with the output of one prompt serving as input for the next. Here, we apply a chain of two prompts where the first prompt handles extraction and its output is the input to the second prompt that handles the pre-processing and structuring the biomarkers in the JSON output.

Swapping GPT-3.5-Turbo with GPT-4 led to better performance with an F2 score of 0.56 for the inclusion biomarkers and 0.42 for the exclusion biomarkers.

NousResearch/Hermes-2-Pro-Mistral-7B (https://huggingface.co/NousResearch/Hermes-2-Pro-Mistral-7B) is a large language model that excels at generating structured JSON outputs. Moreover, the model it was fine-tuned from, Mistral-7B, exhibits superior performance over LLAMA-7B across all benchmarks and outperformed LLAMA-34B in mathematical and coding tasks ^27^.

Hermes-2-Pro-Mistral-7B, demonstrated remarkable capabilities, outperforming the closed-source models at extracting both inclusion and exclusion biomarkers, achieving F2 scores of 0.98 and 0.66, respectively.

Splitting the task into two subtasks with the prompt chaining technique where the first prompt’s output is the input to the second had opposite effects on GPT-3.5-Turbo and GPT-4. For GPT-3.5-Turbo, prompt chaining resulted in a noticeable decline in performance compared to zero-shot prompting. The F2 score for the inclusion biomarkers decreased by approximately 42%, while the F2 score for the exclusion biomarkers decreased by around 16%. In contrast, GPT-4 ’s performance increased with prompt chaining to reach an F2 score of 0.76 for the inclusion biomarkers and 0.70 for the exclusion biomarkers. Few-shot prompting (1S and 2S) showed improvement over zero-shot prompting and prompt chaining for GPT-3.5-Turbo. However, providing two-shot prompting did not improve over one-shot prompting; overall there was a slight decrease in performance. This finding is surprising, as we generally observe an increase in performance with an increase in the number of examples provided to the model ^28^, suggesting that the selected examples might have introduced some conflict or caused the model to overfit.

In Table 2 we show the results with a more mature evaluation approach where we assess the models’ ability to not only extract the biomarkers correctly but to also structure the biomarkers in disjunctive normal form (DNF) in the JSON output as seen in Figure 3. GPT-3.5-Turbo with zero-shot prompting performed the worst with an F2 score of nearly zero for the inclusion and exclusion biomarkers. GPT-4 had an overall moderate performance with an F2 score of 0.29 for the inclusion and an F2 score of 0.43 for the exclusion biomarkers, hence outperforming GPT-3.5-Turbo. The open-source model Hermes-2-Pro-Mistral-7B with zero-shot prompting outperformed GPT-3.5-Turbo and GPT-4. At the extraction of the inclusion biomarker, Hermes-2-Pro-Mistral-7B was approximately 3.24 times better than GPT-4 with an F2 score of 0.94, and around 1.5 times better for the exclusion biomarkers with an F2 score of 0.65.

**Figure 3:**
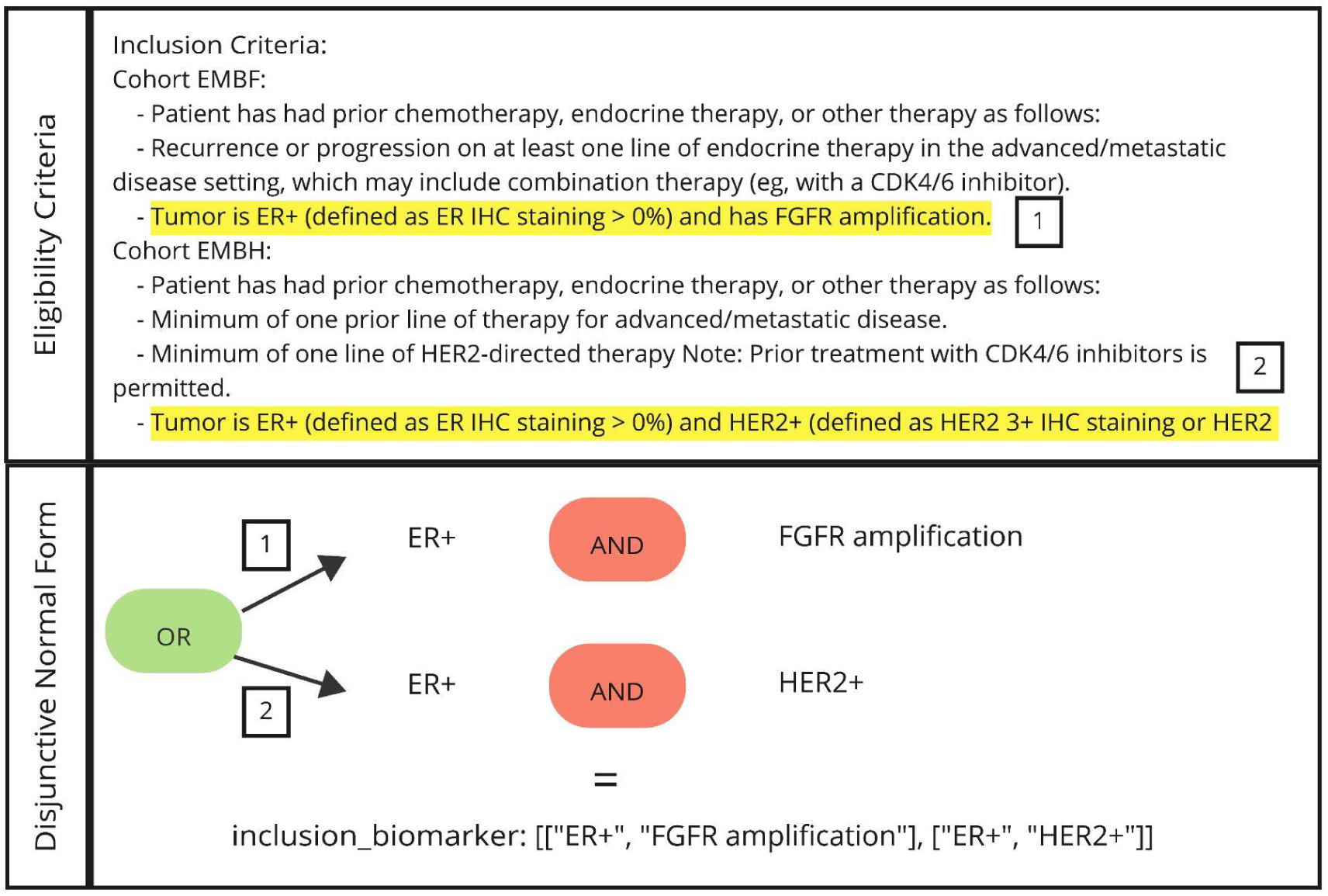
Representation of Extracted Biomarkers in the Disjunctive Normal Form. The figure represents a snippet of the inclusion criteria in the clinical trial NCT04092673. The figure highlights the inclusion requirements for two cohorts EMBF and EMBH. Cohort one EMBF requires the tumor to be ER+ with FGFR amplification. Cohort two required the patient’s tumor to be ER+ and HER2+. These criteria are translated into a structured JSON output in the Disjunctive Normal Form (DNF), reflecting that patients are eligible if they meet either the condition of ER+ with FGFR amplification or ER+ with HER2+.

**Table 2:**
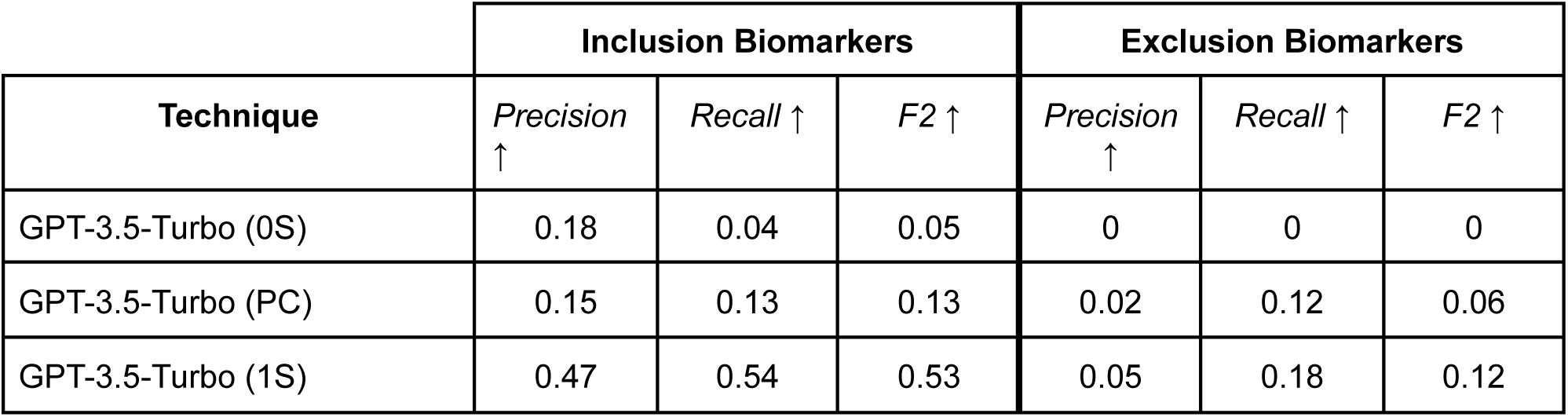
Performance of the open-source and closed-source base models using the different prompting techniques (zero-shot (0S), one-shot (1S), two-shot (2S) and prompt chaining (PC)) in extracting the inclusion and exclusion biomarkers from clinical trials documents while structuring them in the JSON output while adhering to the disjunctive normal form (DNF).

| Technique | Inclusion Biomarkers |  |  | Exclusion Biomarkers |  |  |
| --- | --- | --- | --- | --- | --- | --- |
|  | *Precision* ↑ | *Recall* ↑ | *F2* ↑ | *Precision* ↑ | *Recall* ↑ | *F2* ↑ |
| GPT-3.5-Turbo (0S) | 0.18 | 0.04 | 0.05 | 0 | 0 | 0 |
| GPT-3.5-Turbo (PC) | 0.15 | 0.13 | 0.13 | 0.02 | 0.12 | 0.06 |
| GPT-3.5-Turbo (1S) | 0.47 | 0.54 | 0.53 | 0.05 | 0.18 | 0.12 |

Employing GPT-3.5-Turbo with prompt chaining (PC) increased its performance slightly to reach an F2 of 0.14 for the inclusion biomarkers and an F2 score of 0.06 for the exclusion biomarkers. However, for GPT-4, applying chain prompting led to a decrease in performance, specifically, the model’s F2 score decreased by 7% compared to zero-shot prompting in both inclusion and exclusion biomarker extractions.

Providing an example to demonstrate the task to GPT-3.5-Turbo (1S) caused a surprising improvement in its performance for the inclusion biomarkers to have an F2 score of 0.53. However, for the exclusion biomarkers even though there was an improvement compared to the zero-shot prompting (0S) and prompt chaining, the model still performed poorly with an F2 score of 0.12. When comparing two-shot prompting (2S) to one-shot prompting (1S) for GPT-3.5-Turbo, the F2 score was nearly zero. In contrast, there was no noticeable change in the performance of the inclusion biomarkers.

### Fine-tuned model performance

As mentioned in the previous section, applying Hermes-2-Pro-Mistral-7B with zero-shot prompting already showed impressive results in performing our task, compared to the closed-source models. However, we notice a discrepancy between inclusion and exclusion in the model’s abilities to extract the biomarkers and organize them in the DNF form as seen in table 1 and table 2. To potentially improve the extraction of exclusion biomarkers, we fine-tuned the model using Direct Preference Optimization (DPO). The purpose of fine-tuning is to adapt the LLM to a more specialized task by adjusting its parameters. The DPO algorithm adjusts the model’s parameters to increase the likelihood of generating the desired output by dragging the probability distribution towards the preferred answer and away from the undesired one. Here, we fine-tuned two models initiated from Hermes-2-Pro-Mistral-7B. The first model Hermes-FT and the second Hermes-FT-synth with the labeled datasets DPO-92 and DPO-156, respectively. For more details on the datasets, please refer to the Methods section “DPO Dataset Preparation”. Table 3 demonstrates how fine-tuning the model with DPO-92 which includes 92 manually annotated samples decreased the overall performance. We observe a 10% decrease in the F2 score for the extraction of inclusion biomarkers and approximately a 7.5% decrease for the extraction of exclusion biomarkers. Nevertheless, Hermes-FT-synth, fine-tuned with a larger dataset (DPO-156) demonstrated superb capabilities, outperforming all models with F2 scores of 0.90 and 0.93 for inclusion and exclusion biomarkers, respectively. Despite the overall superiority in performance, the model demonstrated a slight decrease in recall for inclusion biomarkers extraction was observed compared to the base model, Hermes-2-Pro-Mistral-7B. This suggests that the fine-tuned model overlooked some biomarkers during extraction. This behavior could be attributed to the model learning to optimize the extraction of exclusion biomarkers during fine-tuning, thereby reducing hallucination and false positives. It is also possible that during fine-tuning, the model experienced changes in how it handles inclusion and exclusion criteria, causing it to provide brief responses to minimize hallucination in inclusion biomarkers, which led to a decrease in recall.

**Table 3:** Performance of the fine-tuned open-source models with Direct Preference Optimization (DPO) using zero-shot prompting in extracting the inclusion and exclusion biomarkers from clinical trial documents. The models Hermes-FT and Hermes-FT-synth are fine-tuned from Hermes-2-Pro-Mistral-7B with a training dataset of 92 and 156 samples, respectively.

| Technique | Inclusion Biomarkers |  |  | Exclusion Biomarkers |  |  |
| --- | --- | --- | --- | --- | --- | --- |
|  | *Precision*<br>*n* ↑ | *Recall* ↑ | *F2* ↑ | *Precision*<br>*n* ↑ | *Recall* ↑ | *F2* ↑ |
| Hermes-FT (0S) | 0.65 | **0.97** | 0.88 | 0.25 | 0.95 | 0.61 |
| **Hermes-FT-synth (0S)** | **0.99** | 0.88 | **0.90** | **0.82** | **0.96** | **0.93** |

In table 4, using Hermes-FT to extract and structure the biomarkers in DNF yielded unexpected results: a decrease in inclusion biomarkers’ performance with an F2 score of 0.85, alongside a marginal increase in extraction of exclusion biomarkers with an F2 score of 0.67. Hermes-FT-synth resulted in the overall best performance with an F2 score of 0.86 for inclusion biomarkers and 0.94 for exclusion biomarkers.

**Table 4:** Performance of the fine-tuned models Hermes-FT and Hermes-FT-synth using zero-shot prompting in extracting the inclusion and exclusion biomarkers from clinical trials documents while structuring them in the JSON output while adhering to the disjunctive normal form (DNF).

| Technique | Inclusion Biomarkers |  |  | Exclusion Biomarkers |  |  |
| --- | --- | --- | --- | --- | --- | --- |
|  | *Precision* ↑ | *Recall* ↑ | *F2* ↑ | *Precision* ↑ | *Recall* ↑ | *F2* ↑ |
| Hermes-FT (0S) | 0.63 | **0.93** | 0.85 | 0.31 | 0.95 | 0.67 |
| **Hermes-FT-synth (0S)** | **0.99** | 0.84 | **0.86** | **0.86** | **0.96** | **0.94** |

## Discussion

In this study, we explore the capabilities of both closed-source and open-source LLMs to enhance patient matching to biomarker-driven oncology trials. Our results demonstrated a significant discrepancy between the closed-source models and the open-source model. This observed difference suggests that an increase in the volume of the model does not necessarily mean better performance ^29^ and highlights the importance of the training process. The closed-source models GPT-3.5-Turbo and GPT-4 underwent Reinforcement Learning with Human Feedback (RLHF) which helped the models excel at various standard natural language tasks. However, RLHF suffers from a phenomenon known as the “alignment tax” ^30^, which states that the model’s ability to generate more human-like responses is at the expense of the model’s deeper understanding of tasks ^31,32^. In our work, where the model is required to handle a multi-step task, GPT-3.5-Turbo and GPT-4 models struggle to fully comprehend this complex task, even when applied with prompt chaining, one-shot, and two-shot prompting. While the 7 billion parameter model, Hermes-2-Pro-Mistral-7B, that underwent supervised fine-tuning from Mistral-7B, with zero-shot prompting had overall better reasoning capabilities outperforming the closed-source models.

In our work, the open-source model, out-of-the-box, demonstrated superior performance. However, the model presented a disparity in accuracy between the inclusion and exclusion biomarkers. Fine-tuning the model with Direct Preference Optimization using a training dataset consisting of manually samples generated and synthetically generated samples by GPT-4 not only helped close this gap but also achieved the best overall performance (Hermes-FT-synth).

Our study also reveals several key findings. For instance, GPT-3.5-Turbo with few-shot prompting performed competitively with GPT-4 in zero-shot prompting. However, the results highlight the importance of the examples selected as we notice a significant increase in performance when extracting the inclusion biomarkers while the model still suffered at extracting the exclusion biomarkers. Furthermore, with two-shot prompting the model’s performance decreased suggesting a bias in the first example towards our test set since adding a second example introduced conflict for the model causing it to hallucinate more. We discovered that prompt chaining was less reliable than zero-shot prompting. This might be that in prompt chaining the prompts are interdependent leading to a decrease in coherence, and leaving the model with not enough context to efficiently extract biomarkers ^33^. Another key finding is that fine-tuning with DPO presents challenges related to dataset size. Fine-tuning with a relatively small dataset decreased the model’s performance (Hermes-FT) due to overfitting and poor generalization ^34^. Augmenting the training set with synthetically generated samples with GPT-4 improved the model’s (Hermes-FT-synth) overall surpassing other models without having negative effects.

Our results demonstrated that leveraging LLMs to extract a structured output while adhering to the disjunctive normal form is possible. This added layer of complexity provided insights into the models’ reasoning abilities and capacity to handle real-world clinical data effectively. An adequate amount of training data is crucial since it allows for a successful supervised fine-tuning. Augmenting manually annotated samples with synthetic data reduces the time needed for the generation of a fully human-annotated dataset, opening the door for improved fine-tuning.

This study is not without its limitations. The dataset we used for testing our models’ performance is rather small. This small dataset allows us to compare the models’ performance and demonstrate the potential of LLM in performing complex tasks requiring reasoning abilities. However, the test set does not fully encapsulate the complexity of clinical trials criteria. Moreover, in few-shot learning, the selected examples showed a clear bias to the samples in our test set. To avoid such overfitting, more advanced prompting techniques, such as Retrieval-Augmented Generation ^35^, could be applied. While our study incorporated a wide range of prompting techniques and fine-tuning with DPO, two suggestions could help improve performance: hyperparameters tuning during supervised fine-tuning, and testing newly released models such as LLAMA3 (https://github.com/meta-llama/llama3) that may provide better results. In addition, we are only focusing on biomarkers in this study, other criteria such as disease stage, disease type, age, gender, and previous therapies are also important for eligibility for an oncology trial. However, with the increasing importance of biomarkers in oncology therapies the goal of this work is to improve this aspect of trial-matching performance.

## Methods

### Dataset curation and annotation

We retrieved and processed clinical trials from the clinicaltrials.gov database and performed keyword filtering to focus our study on oncology-related clinical trials, which resulted in around 15,856 trials stored in the vector database, ChromaDB. A clinical trial is a text document with a brief description mentioning the purpose of the trial, a detailed description of the trial and the participation criteria that lists the inclusion criteria required for a patient to be considered for enrollment and the exclusion criteria that if the patient possesses would disqualify them from participating in the trial. Our database, ChromaDB, includes only the “brief description” and “eligibility criteria” sections from the clinical trials document.

To select trials that contained genomic biomarkers from our database, we used a list of 500 genomic biomarkers from the CIViC database to perform a semantic search against the cancer clinical trials.

Next, we manually annotated 166 clinical trials by assigning each trial the corresponding JSON output containing the inclusion and exclusion biomarkers present in that trial.

After annotation, we assessed the token distribution to identify outliers that exhibit a significantly larger token count, which we removed to ensure uniformity in the data for downstream analysis. We used 70% of the trials for supervised fine-tuning as a training dataset (train set) and 30% as a test dataset Manual Annotation: The expected output is JSON with two keys, *“inclusion_biomarker”* and *“exclusion_biomarker”*. The *“inclusion_biomarker”* contains the genomic biomarkers required for a patient’s consideration in the enrollment process. In contrast, *“exclusion_biomarker”* contains genomic biomarkers that result in the patient being excluded from the trial enrollment process. To accurately capture the logical connections between biomarkers in the trial (AND/OR logic), we represent them in the Disjunctive Normal Form (DNF).

The DNF which can be described as the disjunction (OR) of conjunctions (ANDs), provides a standardized approach to expressing complex logical connections. In the context of our study, we implement a list of lists data structure to represent the biomarkers in the DNF formula. Each inner list represents a conjunction clause (AND), where all biomarkers conditions in that list must be satisfied. The outer list connects these conjunction clauses through disjunction (OR), where only one of the inner conjunction clauses must be met. This DNF structure allows for a better representation of the complex relationships between genomic biomarkers in clinical trials, enabling more precise patient-trial matching.

### Generation of Synthetic Dataset

To augment the training dataset used in the supervised fine-tuning process, we generated synthetic samples with GPT-4 employed with the prompt provided in Supplementary Fig.2. We included four examples selected from our training dataset in the prompt, with the placeholders in the template indicating where we inserted the examples. Additionally, we appended at the end of the prompt “Trial:” to encourage the language model to generate a new clinical trial that matches the style and format of the given examples. The generated samples comprise a clinical trial input and the corresponding JSON output containing the inclusion and exclusion biomarkers.

Additionally, we predicted the JSON output for our four examples using GPT-4 with zero-shot prompting (Supplementary Fig.3) to ensure the output was coming from the same distribution to increase the consistency and reliability of the generated samples.

Then, we manually reviewed the generated trials to ensure their quality and that they do not contain any unusual or unexpected content that could negatively impact the performance. Finally, we added the synthetically generated samples to the manually annotated training set.

### DPO Dataset Preparation

In this section, we describe the process we followed to prepare our dataset for fine-tuning with Direct Preference Optimization (DPO) ^36^. The DPO algorithm is a technique used to optimize the language model for a downstream task. This algorithm optimized the model by increasing the likelihood of preferred responses (winning completions) and decreasing the likelihood of less preferred responses (losing completions). Refer to “Direct Preference Optimization (DPO) Fine-tuning” section for more details.

We generated two datasets for fine-tuning with DPO, the first deriving from the manually annotated dataset (115 samples) and the second from the mixed dataset (195 samples). A dataset suitable for fine-tuning with DPO should include the input prompts, winning completions (*y_W_*) and losing completions (*y_L_*). The winning completions were either obtained through manual annotation or synthetically generated by GPT-4, with the latter being reviewed and validated by human annotators. To generate the losing completion, we sample from the supervised fine-tuned (SFT) model Hermes-2-Pro-Mistral-7B that we used later for fine-tuning, which we will further describe later.

We used the zero-shot prompt template from Supplementary Fig.4 to generate the prompt for each sample. Once the prompts are prepared, we tokenize them with left padding enabled to ensure that all inputs have the same length.

Using the tokenized prompts, we perform inference with the Hermes-2-Pro-Mistral-7B model, setting the temperature to zero. The generated completion is the losing completion. This inference process was repeated for every sample in each of the datasets.

We further split the samples into 80% as a training set to fine-tune the model and 20% as a validation set to avoid overfitting during fine-tuning.

### Language Models

We investigate the performance of the closed-source models from OpenAI GPT-3.5-turbo and GPT-4, as well as the open-source model NousResearch/Hermes-2-Pro-Mistral-7B.

Even though for previous GPT models details about model architecture and training process were transparent, these details have not been disclosed for GPT-3.5-turbo and GPT4 models. From previous models, we know that GPT models are based on decoder-only transformers, a variant of the original transformer architecture ^31^. It was reported that GPT-3.5-Turbo is the most capable model from the GPT-3.5 series ^32^. In this paper, we use the GPT-3.5-Turbo-0125 and GPT-4 (GPT-4-0613) models.

Additionally, the technical report released by OpenAI ^31^ mentions that GPT-4 outperforms their previous 175B parameter GPT-3 model.

Hermes-2-Pro-Mistral-7B is fine-tuned using teknium/OpenHermes-2.5 (https://huggingface.co/datasets/teknium/OpenHermes-2.5) from Mistral-7B ^27^, a pre-trained generative text model with seven billion (7B) parameters and a context length of 8192.

### Prompting Techniques

Language models are versatile multitask learners ^37^, during training on large and diverse datasets, the pre-trained models learn to perform different tasks. The models’ ability to learn a variety of tasks led to the success of natural language prompting, a method used to condition the model to perform a specific task without updating any of the pre-trained weights ^38^. A prompt is used to provide the model with the necessary information to generate the desired output. Multiple techniques of prompting exist, in this study we employ zero-shot prompting, few-shot prompting and prompt chaining.

- **Zero-shot prompting:** The model is given, at inference time, a prompt that includes the description of the task that the LLM should perform. It usually includes details about the task, the user’s input and the format of the output to be returned by the language model ^39^.
- **Few-shot learning:** Often referred to as in-context learning. Few-shot learning is similar to zero-shot prompting, but in addition, at inference, the model is provided with a set of *k* examples that demonstrate the task ^38^. Each example typically includes an input and the corresponding expected output. Typically, in few-shot learning the model is given *k* examples and then followed by the user’s input.
- **Prompt chaining:** Recent work ^40^ has shown that for complex tasks, decomposing the task into subtasks and chaining together prompts of each subtask can increase output quality. The idea behind prompt chaining is that the output from one prompt is the input of the following prompt, with each sub-task building on the last. This approach should in principle allow the model to tackle complex tasks that may be difficult to accomplish in a single prompt incrementally.

In the zero-shot prompts (Supplementary fig.3-4) we start by giving a summary of our task. In our instructions, we define in detail the biomarkers as any changes at the level of chromosomes, genes, or proteins and the exact JSON output expected including a detailed definition of the DNF logic and how to represent it in the output. One step further, to decrease false positives, we added a description of some common eligibility criteria such as cancer type, pregnancy history, age, gender, etc. and instructed the language model to ignore this information since they do not classify as genetic biomarkers. We also reiterate and mention the important information more than once such as to focus on genomic biomarkers and ignore other details. At the end we describe the processing steps to be performed on the extracted biomarkers. It is noteworthy that while GPT and Hermes-2-Pro-Mistral-7B prompts differ in structure and wording, the fundamental details remain the same.

In our study, we applied two forms of few-shot learning: one-shot prompting, where the model is given only one example (*k* = 1), and two-shot prompting, where the model is given two examples (*k* = 2). The one-shot (Supplementary fig.7) and two-shot (Supplementary fig.8) prompts used with GPT-3.5-Turbo are similar to the zero-shot prompt with the exception of the additional examples consisting of a clinical trial input document and the corresponding JSON output.

For the chain of prompts (Supplementary fig.5-6) used with GPT models, the aim was to reduce the number of tokens in the prompt and simplify the task for the language model by splitting it into two subtasks where the first prompt (Supplementary fig.5) focuses on the task of extracting relevant biomarkers from the clinical trial input and returning a list with AND/OR indicators. While the second prompt’s (Supplementary fig.6) focus is processing the biomarkers and structuring the final JSON output with DNF.

**Figure 4:**
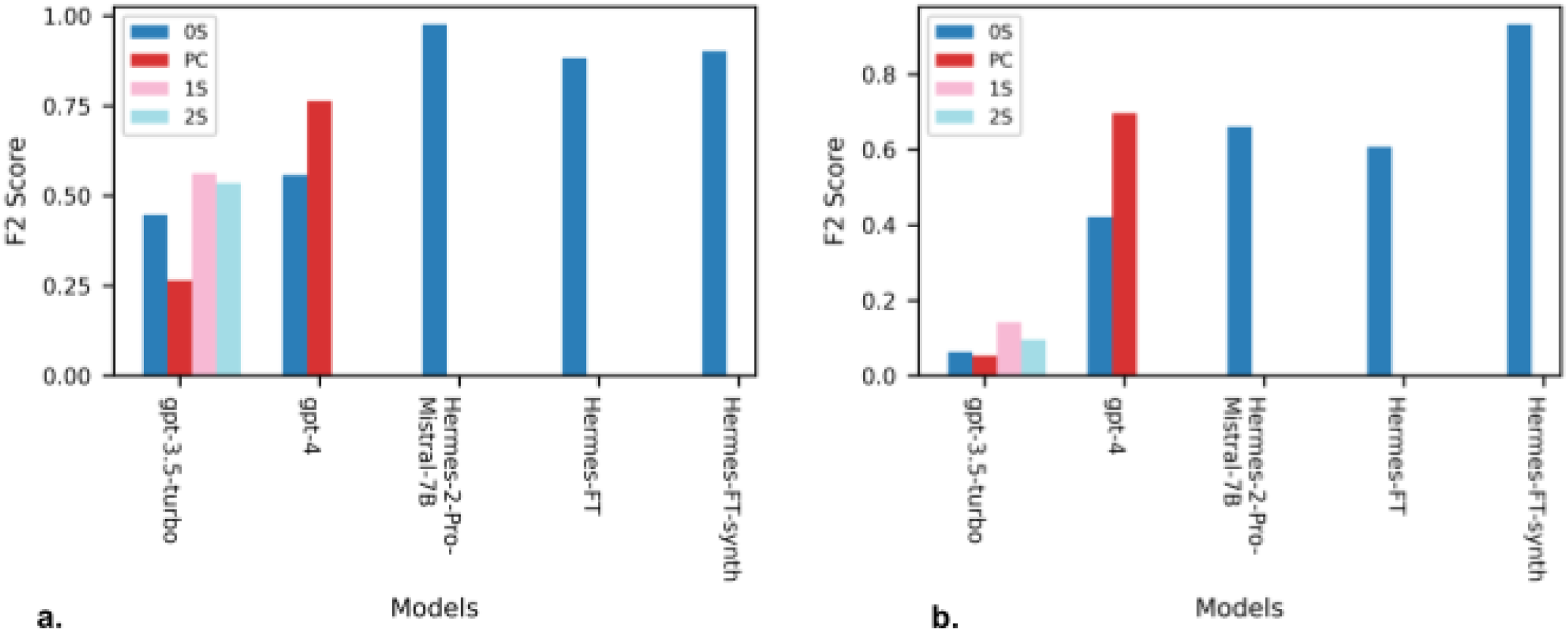
Comparison of biomarker extraction across models. This figure presents a comparative analysis of F2 scores for the base models: gpt-3.5-turbo, gpt-4, Hermes-2-Pro-Mistral-7B, and the fine-tuned models: Hermes-FT and Hermes-FT-synth using multiple prompting techniques: zero-shot (0S), prompt chaining (PC), one-shot (1S), and two-shot (2S). The figure highlights the performance in extracting the inclusion biomarkers (a) and the exclusion biomarkers (b) without considering the model’s abilities in formatting the biomarkers in the Disjunctive Normal Form (DNF).

**Figure 5:**
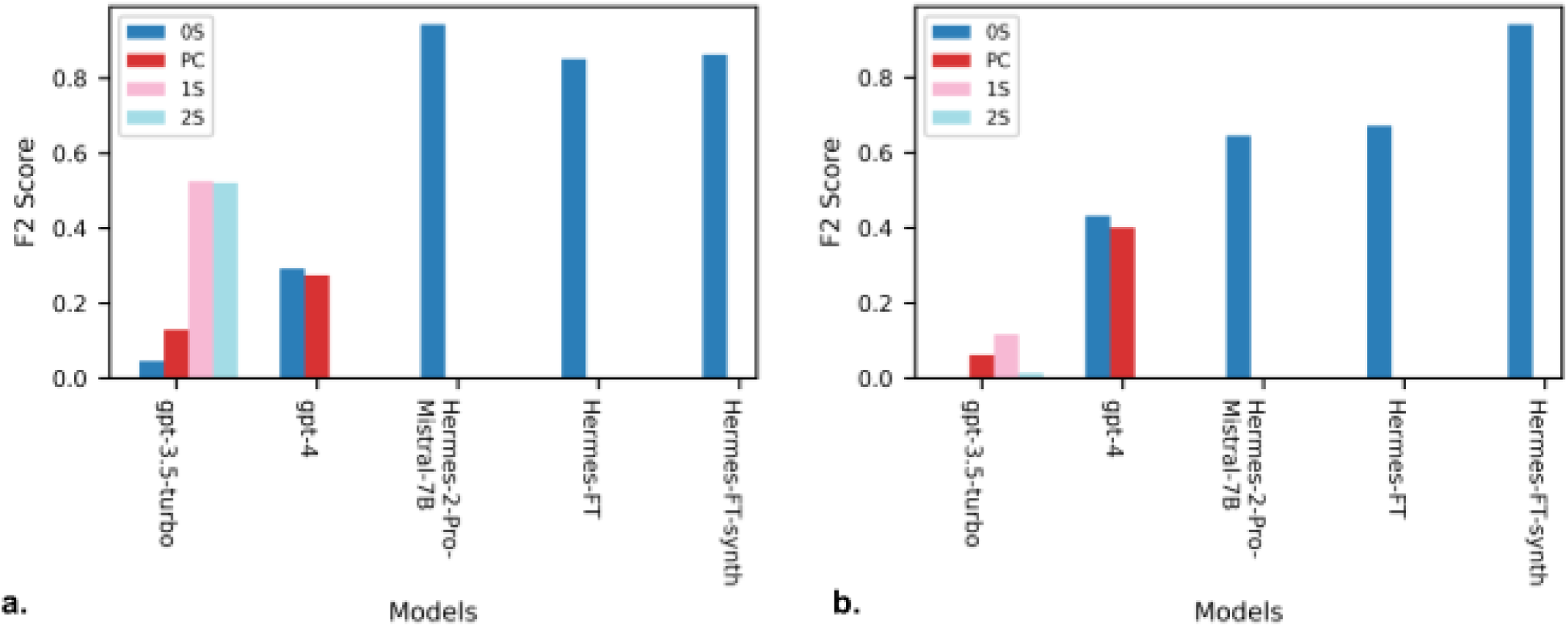
Comparison of biomarker extraction and representation in the Disjunctive Normal Form (DNF) across models. The figure presents a comparative analysis of F2 scores for the base models: gpt-3.5-turbo, gpt-4, Hermes-2-Pro-Mistral-7B, and the fine-tuned models: Hermes-FT and Hermes-FT-synth using multiple prompting techniques: zero-shot (0S), prompt chaining (PC), one-shot (1S), and two-shot (2S). The figure demonstrates the performance in extracting the inclusion biomarkers (a) and the exclusion biomarkers (b) while considering the model’s abilities in organizing the biomarkers in the Disjunctive Normal Form (DNF).

### Direct Preference Optimization (DPO) Fine-tuning

Fine-tuning is a technique used to adapt the LLM to a downstream task and return responses that meet specific requirements. Compared to prompting techniques, during fine-tuning the model’s pre-trained parameters are adjusted. The Direct Preference Optimization algorithm (DPO) optimizes the language model (also referred to as policy) by using preference data directly and minimizing the loss function:

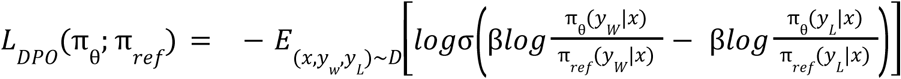

Where *x* is the prompt and *y* is the completion. The annotation *y_W_* refers to the winning completion (desired output), the completion humans prefer over the losing *y_L_* completion (rejected output). This loss function directly optimizes the policy model with parameters θ to increase the probability of the preferred completions π_θ_(*y_W_*|*x*) compared to the dispreferred completions π_θ_(*y_L_*|*x*). The reference model probabilities in the denominator constrain the update to not deviate too far from the original.

Fine-tuning large language models can be very expensive, it requires a lot of GPU memory. Parameter-efficient fine-tuning methods, such as Quantized Low Rank Adaption (QLoRA) ^41^, allow the adaptation of models with a large number of parameters without sacrificing performance by significantly reducing the number of trainable parameters. QLoRA combines the concept of quantization and Low-Rank Adaptation (LoRA) ^42^. The LoRA technique assumes that a small number of weight parameters require adaptation when fine-tuning a pre-trained model with full-rank weight matrix *W*_0_ ɛ *R*^*d*^×*k* for a downstream task. LoRA takes advantage of this by decomposing *W*_0_ into two low-rank matrices *B* ɛ *R*^*d*×*r*^ and *A* ɛ *R*^*r*×*k*^, and rank *r* ≪ *min*(*d*, *k*), such that:

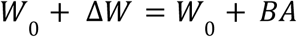

During fine-tuning, *W*_0_ is fixed, and matrices A and B contain the trainable parameters. Initially, *W*_0_ is zero since a random Gaussian initialization is used for *A* and zero initialization for *B*. LoRA also rescales Δ*W* *x* by α/*r*, where α is a constant and *r* is the rank. This low-rank approximation allows efficient fine-tuning of a small subset of adapted weights *B* and *A*. Additionally, QLoRA involves freezing the pre-trained parameters in a quantized representation while fine-tuning the extra set of Low-Rank Adapters.

We employed DPO with QLoRA to fine-tune the Hermes-2-Pro-Mistral-7B model. During the training process, we used the paged AdamW 32bit optimizer with a learning rate of 5e-5, specifying the step size taken by the optimizer during backpropagation to adjust the trainable parameters and minimize the loss function. Moreover, we allow the model to train for 200 steps with a batch size of 8. For the beta factor in the DPO loss function we used the default value 0.1.

For the LORA configuration, the rank is set to 2 and the scaling factor to 4. The trainable parameters are limited to the linear projection layers of the self-attention and multilayer perceptron modules. Furthermore, a dropout probability for the LORA layers of 0.05 is applied.

The Hermes-2-Pro-Mistral-7B model weights are quantized to a 4-bit representation to reduce memory footprint. During the forward and backward propagation, the model weights are represented in Float16, making it suitable for neural network computations while maintaining sufficient precision.

## Data availability

The prompts, data, and results are available at https://github.com/BIMSBbioinfo/oncotrialLLM.

## Code availability

The source code is available at https://github.com/BIMSBbioinfo/oncotrialLLM. Additionally, the fine-tuning and fine-tuned models can be found at https://huggingface.co/nalkhou. Live demo of certain techniques for biomarker-based trial matching can be tested at http://onconaut.ai

## Notes

### Competing Interest Statement

The authors have declared no competing interest.

